# Predicting the effects of epidural stimulation to improve hand function in patients with spinal cord injury: An active learning-based solution using dynamic sample weighting

**DOI:** 10.1101/341719

**Authors:** Mohammad Kachuee, Haydn Hoffman, Lisa D. Moore, Hamidreza Ghasemi Damavandi, Tali Homsey, Babak Moatamed, Anahita Hosseini, Ruyi Huang, James C. Leiter, Majid Sarrafzadeh, Daniel C. Lu

## Abstract

In patients with chronic spinal cord injury (SCI), few therapies are available to improve neurological function. Neuromodulation of the spinal cord with epidural stimulation (EDS) has shown promise enabling the voluntary activation of motor pools caudal to the level of the injury. EDS is performed with multiple electrode arrays in which several stimulation variables such as the frequency, amplitude, and location of the stimulation significantly affect the type and amplitude of motor responses. This paper presents a novel technique to predict the final functionality of a patient with SCI after cervical EDS within a deep learning framework. Additionally, we suggest a committee-based active learning method to reduce the number of clinical experiments required to optimize EDS stimulation variables by exploring the stimulation configuration space more efficiently. We also developed a novel method to dynamically weight the results of different experiments using neural networks to create an optimal estimate of the quantity of interest. The essence of our approach was to use machine learning methods to predict the hand contraction force in a patient with chronic SCI based on different EDS parameters. The accuracy of the prediction of stimulation outcomes was evaluated based on three measurements: mean absolute error, standard deviation, and correlation coefficient. The results show that the proposed method can be used to reliably predict the outcome of cervical EDS on maximum voluntary contraction force of the hand with a prediction error of approximately 15%. This model could allow scientists to establish stimulation parameters more efficiently for SCI patients to produce enhanced motor responses in this novel application.

**Author Summary:** Spinal cord injury (SCI) can lead to permanent sensorimotor deficits that have a major impact on quality of life. In patients with a motor complete injury, there is no therapy available to reliably improve motor function. Recently, neuromodulation of the spinal cord with epidural stimulation (EDS) has allowed patients with motor-complete SCI regain voluntary movement below the level of injury in the cervical and thoracic spine. EDS is performed using multi-electrode arrays placed in the dorsal epidural space spanning several spinal segments. There are numerous stimulation parameters that can be modified to produce different effects on motor function. Previously, defining these parameters was based on observation and empiric testing, which are time-consuming and inefficient processes. There is a need for an automated method to predict motor and sensory function based on a given combination of EDS settings. We developed a novel method to predict the gripping function of a patient with SCI undergoing cervical EDS based on a set of stimulation parameters within a deep learning framework. We also addressed a limiting factor in machine learning methods in EDS, which is a general lack of training measurements for the learning model. We proposed a novel active learning method to minimize the number of training measurements required. The model for predicting responses to EDS could be used by scientists and clinicians to efficiently determine a set of stimulation parameters that produce a desired effect on motor function.

## Introduction

Spinal cord injury (SCI) refers to an acute traumatic injury to the spinal cord that results in varying degrees of sensorimotor deficits below the level of injury [1]. Between 1993 and 2012, the total number of cases of SCI increased in the United States as the population increased, and the total number of new SCIs was approximately 17,000 in 2012 [2]. SCI has a major impact on health-related quality of life and is associated with annual costs of $80,000 per patient after the first year of injury [3]. The acute management of SCI includes surgical decompression and stabilization as well as avoiding secondary injury. The acute management is typically followed by intensive rehabilitation. Despite aggressive intervention for patients with motor complete SCI, which is defined as complete loss of motor function below the level of injury, few patients achieve meaningful neurological recovery with rehabilitation [4].

Among novel interventions for chronic SCI, neuromodulation using epidural stimulation (EDS) caudal to the level of injury has shown promise in allowing patients to regain voluntary locomotor activity [5]. Edgerton et al. discovered that EDS enabled patients with motor and sensory complete SCI to regain fine voluntary movement below the level of injury [6, 7]. In the presence of EDS, patients were able to voluntarily control the force generated in specific muscle groups in response to visual and auditory cues. After repeated training, stimulation thresholds to produce voluntary motor activity decreased. Recently, EDS has been applied to the cervical spine to improve hand function, which is less related to central pattern generation than locomotion. These patients demonstrated up to a 300% increase in hand strength [8].

The foregoing studies used multi-electrode epidural stimulation arrays spanning 2-3 cervical or lumbosacral spinal segments. There are multiple stimulation variables that can be modified to change the effect of EDS, including stimulation amplitude, electrode polarity, stimulation frequency, pulse width, and stimulation location. Optimizing the stimulation configuration and location markedly affects the type and amplitude of efferent motor responses [6, 9, 10]. Identifying the optimal configuration is conventionally performed by empirical methods that systematically explore the array space, which is a time consuming and expensive process. Recently, an active machine learning technique using a structured Gaussian process was used to optimize stimulus variables in four spinally transected rats with implanted multi-electrode epidural arrays [11]. The algorithm was able to select stimuli that produced the best motor response out of a very large set of parameters and independently identify stimulation locations associated with specific tasks. Ultimately, it was able to match the efficacy of human experimenters in identifying the optimal stimulation variables.

Given the extensive input and output information associated with modifying stimuli and characterizing motor responses, respectively, there is a need for efficient machine learning algorithms to explore the wide range of stimulation properties and locations available with EDS. We developed a novel approach for predicting the effect of different stimulation configurations on hand function in a subject with chronic SCI following placement of a cervical epidural spinal cord stimulator. We focused on predicting the outcome of different stimulation configurations in an automated fashion using an active learning method for guiding the experiments. Apart from the prediction of the outcome of stimulation, we also developed a novel method based on artificial neural networks to interpret the outcomes of several experiments and aggregate them as a single target value. This was used to combine EMG signals from different trials into one value. We refer to this process in the following sections as the “dynamic weighting method.”

## Results

Our dataset consisted of 237 samples taken from a single patient with chronic SCI in which each sample was a combination of stimulation parameters (i.e., frequency, intensity, and location) and a target outcome score. In order to evaluate the proposed method, a 10-fold cross-validation of test and training data was used (i.e., training on 9 folds, and testing on the remaining 1 fold while cycling the test fold and averaging all test performance results). Using this strategy, the following results were obtained from all 237 samples.

### Outcome prediction performance

Table 1 presents a comparison between the performance of the proposed dynamic weight prediction method and other simple, commonly used approaches to characterize the outcome such as mean and median. The comparison is made in terms of mean absolute error (MAE), standard deviation (STD) of prediction errors, and Pearson’s correlation coefficient (*r*) between the actual target and the predicted values. As shown in the table, the dynamically weighted predictions outperform the other methods by a considerable margin; the MAE and STD values for the dynamic weighting method are about 27% and 24% lower than the MAE and STD values for the mean and median estimates, and the correlation coefficient is greater. In addition, Table 1 compares the results using i) a neural network predictor trained alongside the dynamic weight predictor, and ii) a support vector regression (SVR) predictor trained on the dynamically weighted targets. From this comparison, training a new SVR predictor instead of using the existing neural predictor yielded more accurate predictions.

**Table 1:** Comparison between the proposed dynamic weighting score weighting approach and mean/median approaches.

|  | **Mean** | **Median** | **Proposed approach (NeuralNet)** | **Proposed approach (SVR)** |
| --- | --- | --- | --- | --- |
| MAE (%) | 21.07 | 21.63 | 20.01 | 15.23 |
| STD (%) | 26.65 | 27.38 | 26.12 | 20.25 |
| $r$ | 0.55 | 0.57 | 0.60 | 0.64 |

MAE, mean absolute error; NeuralNet, neural network; STD, standard deviation of error; SVR, support vector regression.

Fig 1 shows the histogram of errors between target values and predictions using the proposed dynamic weighting approach combined with SVR and the equally weighted averaging approach combined with SVR. The error histogram corresponding to the dynamic weighting approach is more concentrated around zero (i.e., has a lower variance). Moreover, the range of errors associated with the combined dynamic weighting and SVR is smaller than the range of errors associated with equally weighted averages.

**Fig 1.**
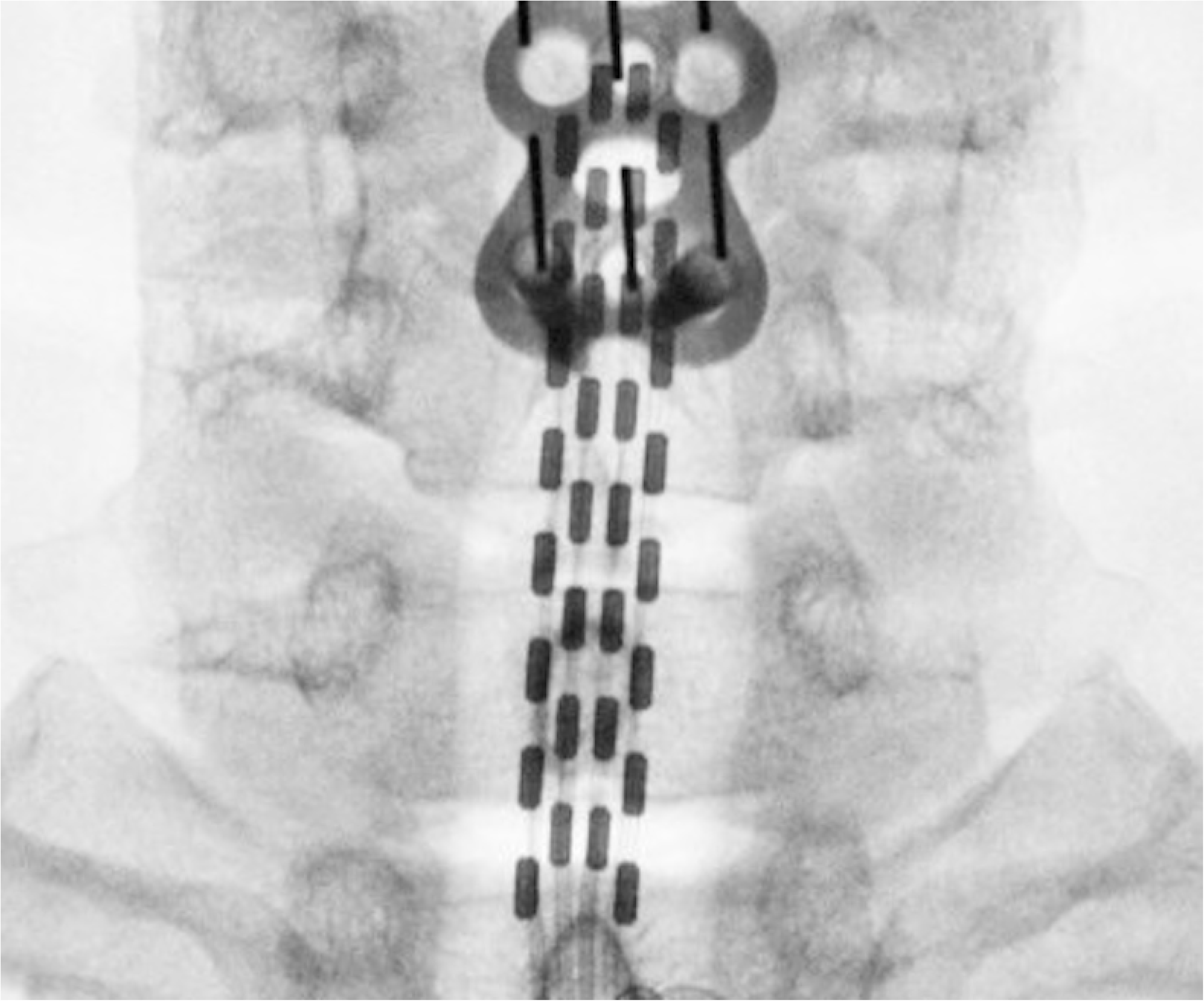
Histogram of errors using dynamic weighting and SVR compared to mean and SVR. Increasing the efficiency of learning.

Fig 2 compares the learning curves derived from using the committee-based active learning method and the random sample selection approaches for *r*, MAE, and STD performance measures. The horizontal axis in these figures is the fraction of the patient’s data from the entire dataset that was used for training the learning methods. The active learning method was able to reach the same accuracy as the random sampling method, as reflected by the r-values, despite using 30% less of training data (Fig 2a). In other words, the active learning method reduced the number of samples required to train the model by approximately 30% to achieve the same prediction performance as the random sample selection. The performance of the active learning approach was superior to the random sampling method, and this improved performance is reflected in the MAE and STD measures seen in Fig 2b and Fig 2c.

**Fig 2.**
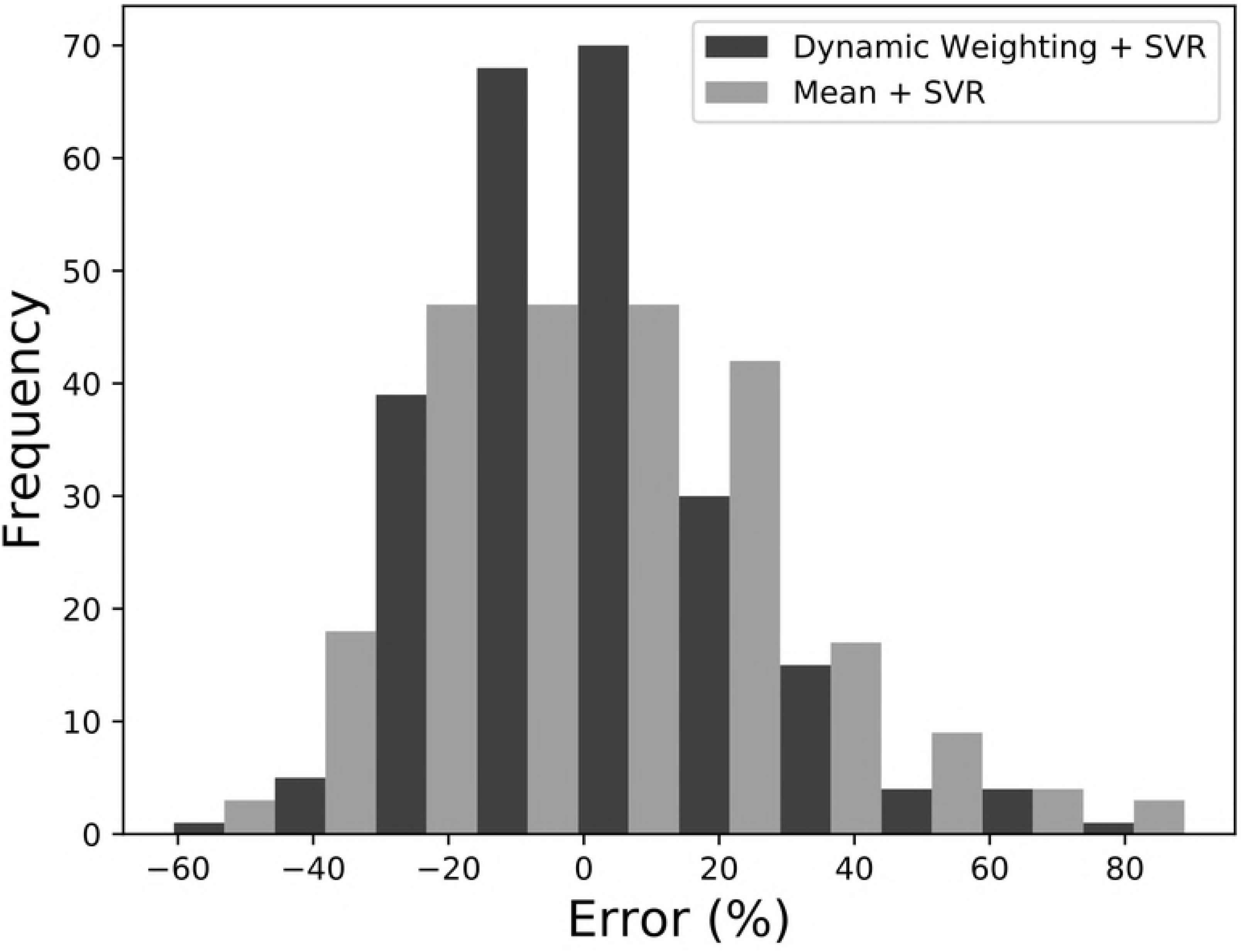
The performance of active learning compared to random sampling in terms of (a) r-value, (b) mean absolute error (MAE), and (c) standard deviation (STD).

In this figure, the X-axis represents the fraction of training data used from the entire available data samples, and the Y-axis represents each performance measure.

## Discussion

Spinal EDS is emerging as a potential therapy for patients with motor complete cervical and thoracic SCI to achieve previously inaccessible purposeful, voluntary movements in the hands and legs. The mechanism by which this occurs is unclear. EDS may directly activate proprioceptive fibers in the dorsal column and dorsal roots [12] or indirectly affect interneurons [13]. As demonstrated previously, different lower extremity movements such as stepping, standing, and isolated motions are subject to substantial variability based on the configuration and location of the electrode array as well as the stimulation parameters [14-17] and may depend on the individual characteristics of each patient. Similarly, the effect of EDS on upper extremity movements is affected by the stimulation parameters applied in the cervical spine [8]. Previously, identifying stimulation parameters has been based on empirical testing and visual observation. Given the inefficiency and time consuming nature of empirically testing every combination of these variables, computational models have been pursued to predict the motor responses produced by EDS based on the intensity and location of the stimulus [18]. We addressed the inefficiency of prior approaches by developing a novel technique using machine learning methods to predict the hand contraction force of a patient with chronic SCI based on different EDS parameters. The accurate prediction of hand contraction force will reduce the lengthy testing sessions needed to obtain stimulation settings empirically. This is the first study to use such an approach in a patient with cervical SCI so far was we know.

In this study, the proposed dynamic weighting method was used to derive a target value of hand force generation for the purpose of training a model to predict the outcome of cervical EDS. Dynamic weighting proved to be more accurate than using equally weighted averages. In general, the dynamic weighting method can be applied to derive a single quantity when there are several measurement samples available, which is useful when the traditional filtering and averaging methods are not sufficient. We limited the active learning queries to a pool of samples that were generated during subject testing before – this was not an exhaustive set of all possible stimulation combinations, but did represent the available dataset from this patient. However, the algorithm could be allowed in future applications to query the entire space of stimulation variables that is reasonable to obtain based on hardware limitations and patient comfort. The committee-based active learning method that we described increased learning efficiency, which can reduce the number of clinical experiments necessary to identify optimal stimulation configurations.

The SVR model produced more accurate predictions of motor function than a neural network. However, it should be mentioned that in other possible applications of the proposed dynamic weighting method, using the jointly trained neural predictor with enough training data might be a reasonable option. In order to limit the number of experiments required to obtain training data, an active learning method similar to the one that we described can be used. This approach used 30% less data than the random sampling method and yielded similar accuracy, which may translate into a genuine saving of time and discomfort for each patient.

The current study has some limitations. Training the model was limited to 237 samples from the subject. Using a larger number of samples would have improved the training of our deep learning model and made it more generalizable. However, given that this is a novel application and with only two cervical SCI patients implanted to date that we are aware of, this study may be timely for future implantations. In this study, we only addressed hand contraction because we had a reliable method for quantifying this. There are a variety of other movements necessary to achieve meaningful hand function, however, and simultaneously optimizing these and other outcome variables during EDS treatment may require modification of our technique. Our approach can likely be applied in a similar manner to the lumbosacral spine as well in order to predict walking ability in response to EDS.

## Materials and methods

### Ethics statement

The study and the experimental protocols were approved by the UCLA Institutional Review Board (IRB#12-001416) and FDA IDE (G140103).

### Experimental setup and data collection

#### Data collection procedures

A 28-year-old male with chronic SCI was enrolled in the current study. The patient sustained an injury at C5 from a motorcycle accident and the severity of his injury was assessed using the International Standards for Neurological Classification of Spinal Cord Injury (ISNCSCI) as an American Spinal Injury Association (ASIA) Impairment Scale (AIS) C.

A 32-contact paddle (Coverage X32, Boston Scientific Corporation) was implanted in the dorsal aspect of the cervical spine (Fig 3). In this study, the paddle contacts in each row were stimulated together. We explored stimulation frequencies of 5, 30, 60 and 90 Hz and intensities between 1 mA to 6 mA.

**Fig 3.**
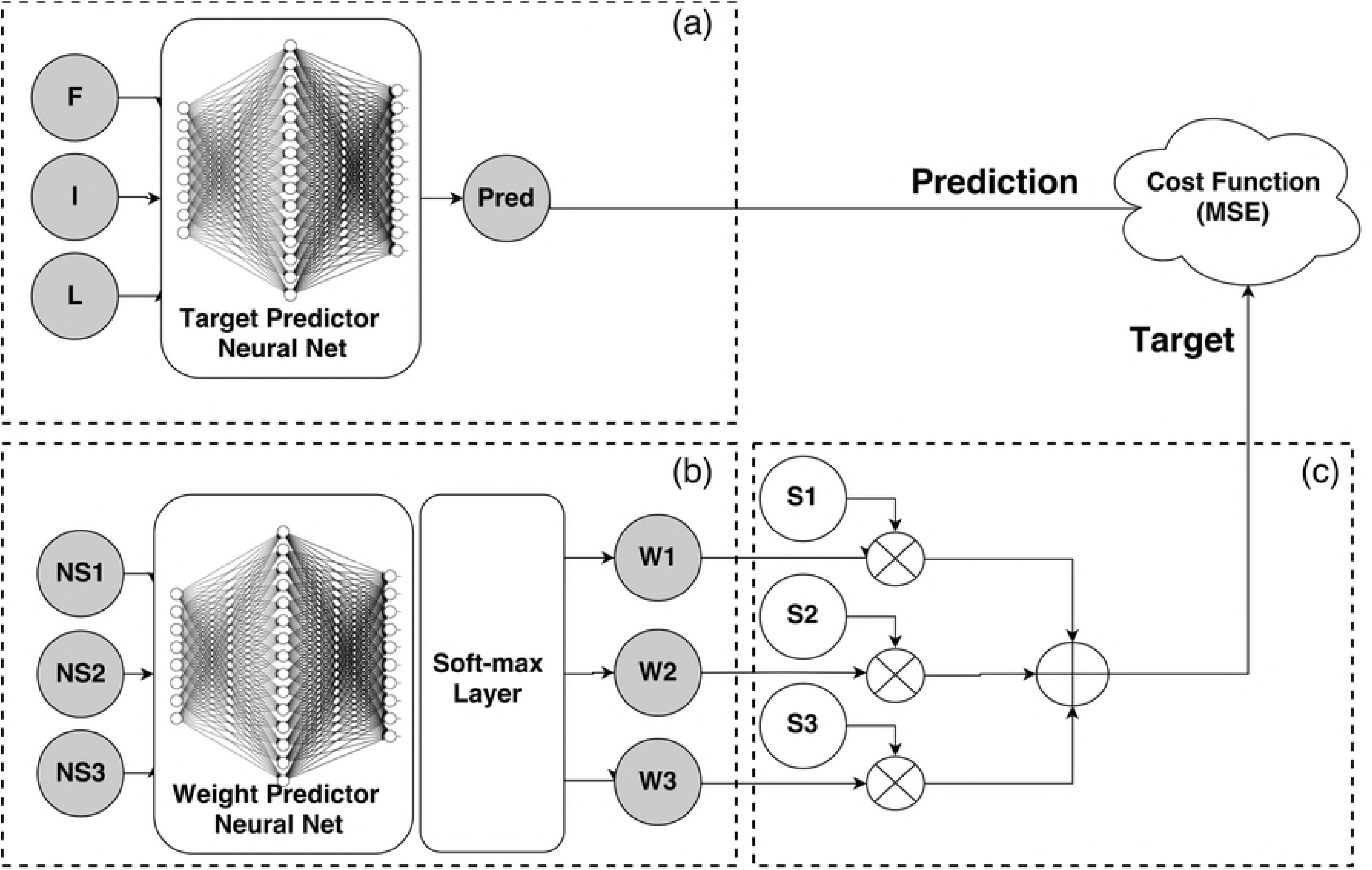
X-ray showing the epidural stimulation paddle contacts and their placement implanted in the cervical spine.

The data used in this study consisted of 29 different experimental sessions (each session on a separate day) that were conducted over a 15-week period. Prior to each session, experimental configurations (i.e., frequency, intensity, and location) were selected based on decisions by a human expert.

#### Hardware setup

A custom designed hand-grip device with adjustable tension control was used for the maximum voluntary contraction (MVC) hand-grip task. The application of this device for quantifying voluntary upper extremity motor function in patients with SCI has been described previously [19]. The tension of the hand-grip device was adjusted to accommodate the patient’s range of potential generated force. A data acquisition device (RZ5D BioAmp Processor, TDT Corporation) was used to record 16 EMG channels as well as a manual synchronization pulse at a sampling frequency of 24.4 KHz. The brachioradialis EMG channel was included in the analysis. Apart from the EMG signal a manual pulse signal, which is henceforth called the synchronization signal, was created manually during the experiments. This was used to mark the beginning and the end of each hand-grip MVC task.

### Study design

This study consisted of 29 different sessions. In each session, the subject was asked to perform multiple MVC experiments, which included baseline (without any stimulation) and during epidural stimulation (using different stimulation configurations). In each experiment, the subject performed three MVC hand-grip trials sequentially (all with the same stimulation configuration). Each trial was initiated with a verbal cue indicating that the subject should grip the device handle as strongly as he could. Each EMG signal interval that corresponded to a single trial within an experiment was called a signal portion. Raw signal portions were used to measure the score of each trial. The score was defined as the median of the top 5% of the highest EMG amplitudes (Fig 4), which was normalized by the baseline score using equation 1. The target score was derived by aggregating the scores of the three trials using mean, median, or the proposed weighting method. A feature vector, a combination of location, intensity, and frequency, defined the stimulation configuration during each particular experiment.

**Fig 4.**
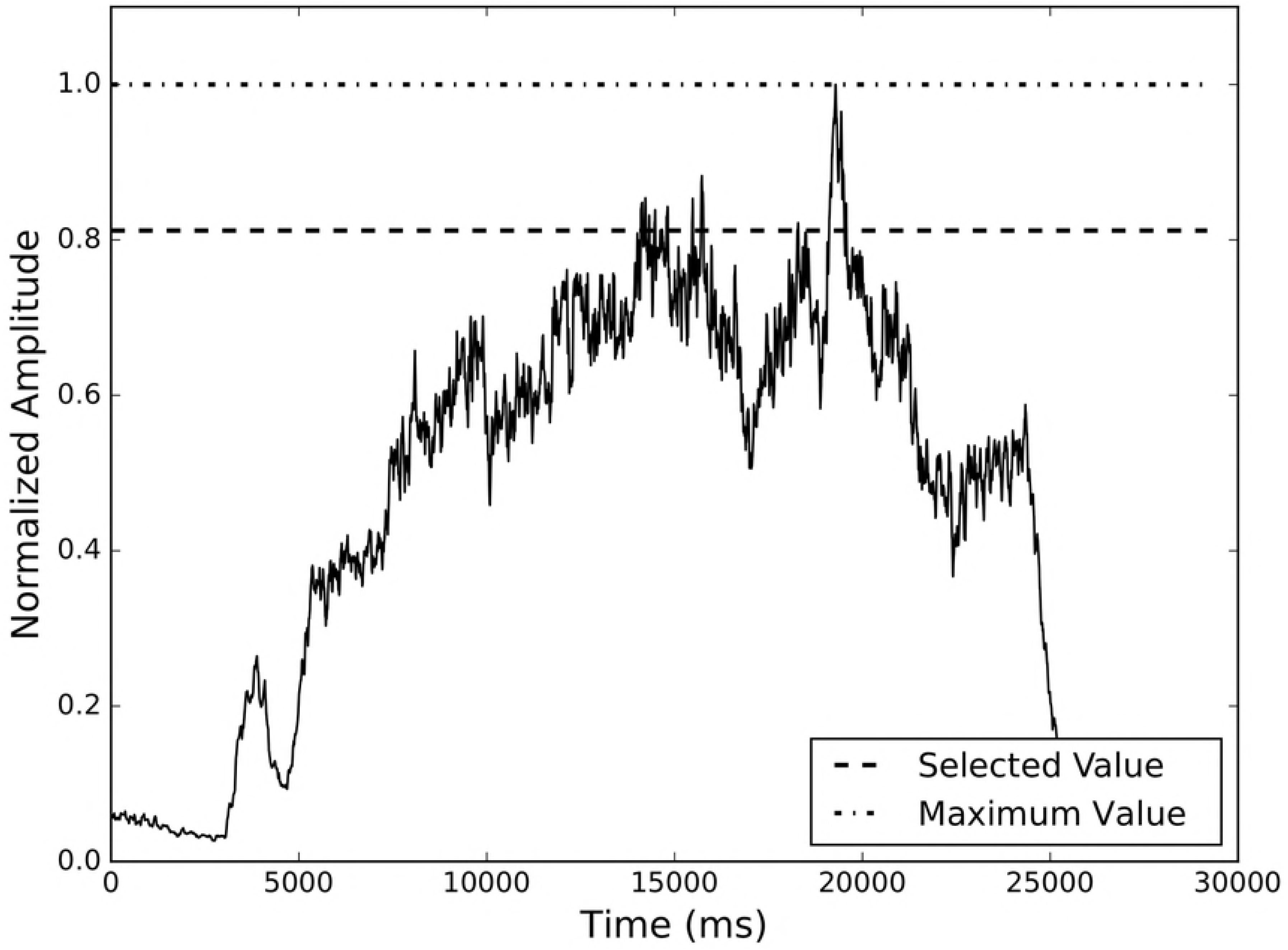
Score calculation in a single EMG portion.

The markers show that it is unreliable simply to use the peak value, and the results are improved using the proposed method.

### Preprocessing

Fig 5 shows the proposed preprocessing steps. Preprocessing started with a 10x decimation (down-sampling by a factor of ten) and full-wave rectification of the EMG and synchronization signals (the original signal had a sampling frequency of 24.4 KHz). The rectified signals were filtered using median filtering with a window size of 200 ms for the synchronization signal and mean filtering with the same window size for the EMG signal to eliminate noise and spike-like values due to analog to digital conversion errors. Next, the EMG and synchronization signals were normalized using unity-based normalization (i.e., scaling values to the range of zero and one). The synchronization signal was used to split the EMG signal to three portions, each corresponding to the EMG signal during a single trial (Fig 6). For each portion, the median amplitude among the 5% of maximum amplitudes was calculated as the score of that trial. The logic behind selecting the median amplitude among the 5% of maximum amplitudes, instead of simply using the peak value, is that the median value significantly increases the robustness of score calculation in the presence of noise and unwanted effects on the EMG signal. The maximum value may be an outlier and not be a good representative of the ‘true’ maximum value of the signal. As shown in Fig 4, simply using the peak value results in inaccurate score measurements due to the noise and artifacts affecting the EMG signal.

**Fig 5.**
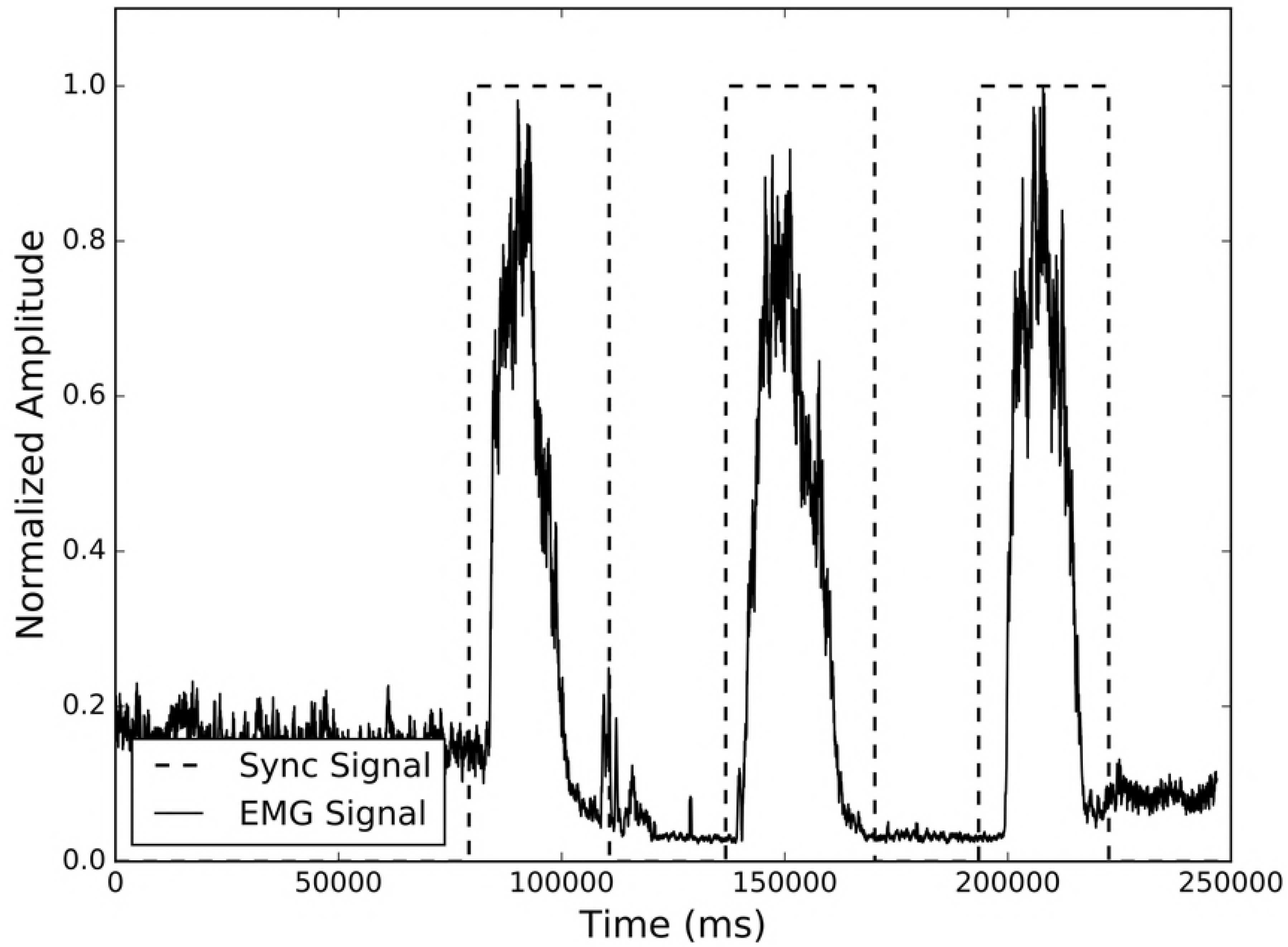
The proposed preprocessing pipeline.

**Fig 6.**
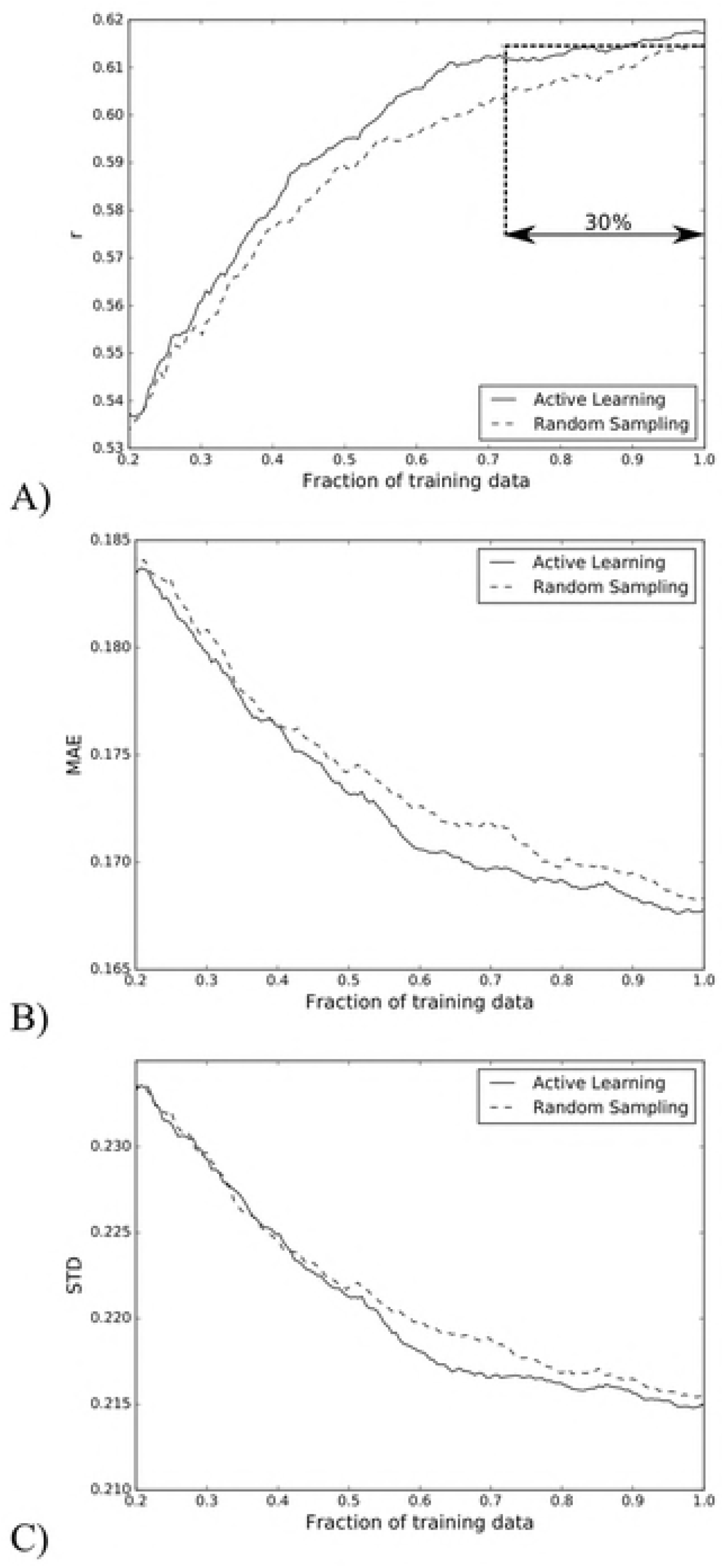
The synchronization signal was used to divide the EMG signal to three portions and each corresponded to a trial within the experiment.

After calculating the scores associated with each experiment within a session, the final scores for each stimulation experiment were calculated using the following formula.
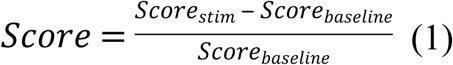

Where *Score_baseline_* is the score before applying any stimulation, and *Score_stim_* is the score during stimulation. The final score is the percentage of variation between the stimulation score and the baseline score obtained during each test session.

### Calculation of target values

Three scores were available for each experiment corresponding to each trial. We calculated a single target score (the aggregated value using a weighted average of the scores for each signal portion where a score for each signal portion is the median of the top 5% of the highest EMG amplitudes) from these scores as the outcome of each experiment. The simplest approach would have been to average these values [20]; however, many scores were contaminated by noise and some could have been limited by subject effort. Accordingly, using a simple average or median of scores may not be an accurate measure of the real expected target values. The problem becomes more complicated considering that, within each experiment, the subject was less fatigued in the first trial compared to the following two. Therefore, we used a weighted average to calculate the target values, and the weights were determined using a predictor model that dynamically predicted the appropriate weights for each trial score. We created two neural networks in which one network was used as an outcome predictor, and the other one was used as a weight predictor, and we trained them jointly. The first neural network was responsible for predicting the stimulation outcome, and the second one was responsible for predicting the weights that were applied to each trial score in the calculation of the target value. Fig 7 demonstrates the proposed network architecture. The dotted areas (a) and (b) in Fig 7 show the outcome predictor and the dynamic weight predictor, respectively.

**Fig 7.**
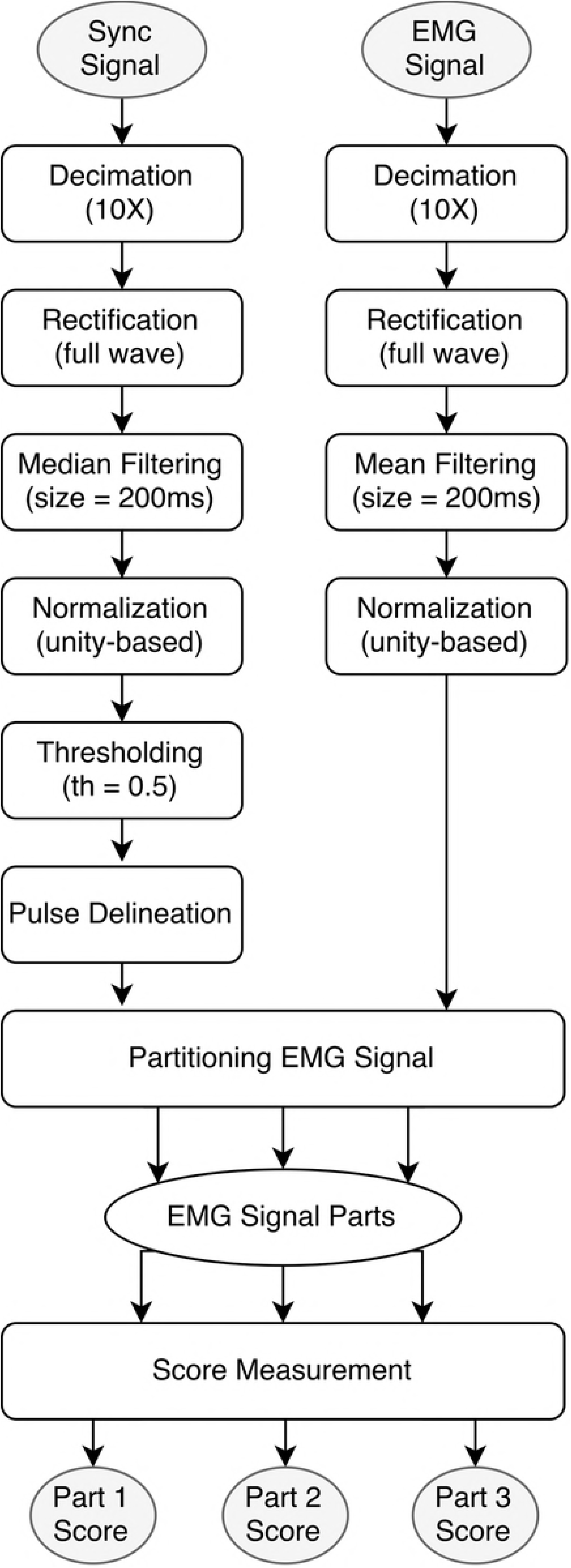
The proposed neural network architecture.

The network can be separated into (a) the outcome predictor, (b) the dynamic weight predictor, and (c) the weighted average calculator. In this figure, F, I, and L correspond to the stimulation parameters frequency, intensity, and location, respectively. NS1, NS2 and NS3 are the three recorded scores (the median of the top 5% of the highest of EMG amplitudes) normalized by their mean value for a given stimulation configuration. Target value is an aggregated value of these scores (not normalized) using the proposed dynamic weighted average of the scores.

In order to direct the network to learn from relative sample values rather than absolute values, the score values normalized by their median values (NS1, NS2, and NS3 in Fig 7) were used as inputs for the weight predictor. The last layer of the weight predictor is a softmax layer, which guarantees that the predicted weights are always positive and sum to one.

After multiplying the score values (the median of the top 5% of the highest EMG amplitudes) with these predicted weights, we estimated the target value, which is the weighted average of the scores (see the dotted part (c) of Fig 7). The softmax layer played a significant role in the training of the whole network and ensured that the weights are positive and sum to one. In the absence of this softmax layer, the network would have chosen an arbitrary set of weights leading to an incorrect target value not indicating the weighted average of the scores. To train the network we used TensorFlow, an open-source numerical computation library developed for dataflow programming that is commonly used to train neural networks [21]. The proposed dynamic weight predictor can theoretically work for any kind of neural network including deep learning methods [22]. However, due to the limited number of training samples available to us, we used networks with a single hidden layer consisting of 32 neurons. The optimization cost function was defined as the mean squared error (MSE) between predictions and targets of the complete network (Fig 5). Finally, stochastic gradient descent (SGD) was used to train the network [23].

### Prediction of target values

The trained outcome predictor in the previous section can be used directly to predict the expected target value of each stimulation configuration. However, from the trained network, we only used the dynamic weight predictor for the estimation of the expected target values. Based on these target values, a Radial basis function (RBF)-kernel support vector regression (SVR) was trained as the final stimulation outcome predictor. We used Scikit-Learn library [24] to train the RBF-kernel ϵSVR model. Model hyper-parameters including the error term penalty (C), error insensitive zone width (ϵ), and kernel smoothness parameter (γ) were selected based on a 10-fold exhaustive grid search cross validation (C = 15; ϵ = 0.01; γ = 0.001).

### Increasing the Learning Efficiency

Active learning methods attempt to decrease the number of labeled training data required by requesting samples that are considered to be more informative. There are a variety of strategies to optimize the selection of informative samples, such as maximum uncertainty sampling, query by committee, expected model change, and expected error reduction [25]. Active learning classification and clustering methods have been widely studied [25-27], but only a limited number of these methods address regression problems [28, 29].

Due to the considerable time, cost, and effort required to collect clinical data, an active learning method may guide data collection and the exploration of stimulation space efficiently and at less cost. In the current study, however, we had already collected a reasonable number of samples, so we simulated the active learning by starting with a small, randomly selected portion from all training samples and considered the remaining samples unlabeled. In other words, we created an unlabeled pool (while we actually knew the true labels), and only included these samples in the training once queried by the active learner. A committee of eight SVR models was trained using bootstrap aggregating, each time selecting 90% of training data randomly with replacement. Afterwards, the variance of committee member predictions was used to find samples (or, configurations) with the highest disagreement or equivalently highest variance values. Finally, the sample with the highest disagreement was selected as the next sample to be explored.

## Acknowledgements

This research was made possible by generous support from the H&H Evergreen Foundation and the NIH: EB15521 funded by NIBIB, NINDS, and NICHD. The research described was conducted in the UCLA Clinical and Translational Research Center (CTRC) which was supported by NIH/National Center for Advancing Translational Science (NCATS) UCLA CTSI Grant Number UL1TR000124. D.C.L. is a 1999 Paul Daisy Soros New American Fellow.

